# Viruses Diversity and Interactions with Hosts in Deep-Sea Hydrothermal Vents

**DOI:** 10.1101/2022.08.13.503714

**Authors:** Ruolin Cheng, Xiaofeng Li, Lijing Jiang, Linfeng Gong, Claire Geslin, Zongze Shao

## Abstract

**Background:** Deep-sea harbor enormous viruses, yet their diversity and interactions with hosts in hydrothermal ecosystem are largely unknown. Here, we analyzed the viral composition, distribution, host preference and metabolic potential in different inhabits of global hydrothermal vents.

**Results:** From 34 samples of eight vent sites, a total of 4,662 viral populations were recovered from the metagenome assemblies, encompassing diverse phylogenetic groups and defining many novel lineages. Apart for the abundant unclassified viruses, tailed phages are most predominant across the global hydrothermal vents, while single-stranded DNA viruses including *Microviridae* and small eukaryotic viruses also constitute a significant part of the viromes. These viral populations were grouped into 1,138 genus-level clusters by protein-sharing network analysis. More than half were exclusively of hydrothermal origin, reflecting the remarkable novelty of hydrothermal viruses. Among the typical niches, vent plumes own the largest number of viral clusters compared to diffuse flows and sediments. Moreover, merely 11% of the viral populations can be linked to specific hosts, which are the key microbial taxa of hydrothermal habitats, such as Gammaproteobacteria and Campylobacteraeota. Intriguingly, vent viromes shared some metabolic features in common that they encode auxiliary genes extensively involved in metabolisms of carbohydrate, amino acid, cofactors and vitamins. Specifically in plume viruses, various auxiliary genes related with the methane, nitrogen and sulfur metabolisms were observed, indicating their contribution to host’s energy conservation. Moreover, the prevalence of sulfur relay pathway genes notified the significant role of vent virus in stabilizing tRNA structure, which promotes host’s adaptation to the steep environmental gradients.

**Conclusions:** The deep-sea hydrothermal systems reserve an untapped viral diversity with novelty. They may affect both vent prokaryotic and eukaryotic communities, and modulate host metabolisms related to vent adaptability. More explorations are needed to depict global vent virus diversity and their role in the unique ecosystem.

## Background

Deep-sea hydrothermal vents are one of the most extreme and dynamic environments on Earth [1]. In this dark world, mixing between the anoxic, hydrothermal fluids and oxic, cold seawater results in wide chemical and thermal gradients, providing energy sources for the vent ecosystems [2]. Unlike most ecosystems that are fueled by photosynthesis, biological productivity is primarily driven by chemoautotrophs in deep-sea hydrothermal plumes [3], the chemoautotrophs use the energy produced by oxidation of sulfur, hydrogen, methane, ammonia or iron to fix carbon [2, 4], converting dissolved inorganic carbon into the organic phase within the biota. The hydrothermal plumes can rise hundreds of meters and disperse hundreds of kilometers away from their source, and impact on broader deep-sea microbial communities and biogeochemistry [5]. In the past decade, significant efforts have been made to explore the source, diversity and function of the microbes inhabiting the hydrothermal plumes [3, 6–10]. These studies suggested that the prokaryotic communities in hydrothermal plumes are distinct from those in the diffuse fluids [11], and those of the subseafloor habitats [12, 13]. The viral communities, as important components of the hydrothermal vent microbiomes, however, have received less attention.

Viruses are the most abundant, pervasive and genetically diverse biological entities of the biosphere [14]. In the ocean, the total estimated numbers of viruses are about 10^30^, composing the second largest relative biomass (but the most abundant) by comparison with prokaryotes and protists, despite their small size [15, 16]. They play a pivotal role in marine ecosystems not only through lysing their hosts, but also through horizontal gene transfer and manipulation of host metabolism [17]. Each day, viruses in surface waters kill 20-40% of the prokaryotes and release up to 10^9^ tons of carbon and other nutrients, which has a significant influence on ocean biogeochemical cycles [16]. Meanwhile, it is estimated that marine viruses transduce approximately 10^14^–10^17^ Gbp of DNA per day [18], affecting host diversity and function. Comparatively, our knowledge of viral diversity and processes in the deep sea is quite limited, partially due to the difficulties in obtaining and processing samples of deep sea.

In the deep-sea hydrothermal vent ecosystems, virus-like particles are more abundant than prokaryotes, and are believed to have a profound impact on microbial communities [19]. The viral abundances of hydrothermal plume samples were reported to be 10^5^–10^6^ VLPs ml^−1^, higher than the surrounding seawater samples [20, 21]. Moreover, it has been suggested that the hydrothermal vent microbes harbor substantial populations of temperate viruses [22–24], which may improve host fitness and facilitate horizontal gene transfer. A well-known example of phage auxiliary metabolic genes (AMGs) is the gene encoding the reverse dissimilatory sulfite reductase (*rdsr)* [25]. This enzyme was identified in hydrothermal plume phages that putatively infect sulfur-oxidizing bacteria, suggesting that viruses play a direct role in the sulfur cycle. Besides, viral AMGs are involved in various metabolic pathways including nitrogen, methane metabolism but also in DNA replication/repair/recombination and amino acid biosynthesis [24, 26], or even compensate novel metabolic pathways for their host microorganisms [19]. These findings suggest that hydrothermal vent viruses are a large reservoir of genetic diversity and have complex interactions with their hosts and habitats, which remain to be fully elucidated.

So far, microbes identified in hydrothermal vent habitats are largely uncultured and few virus isolates have been reported [27, 28]. Thus, metagenomics technology is commonly applied to characterize the diversity, ecology and evolution of hydrothermal vent viruses [19, 22–25, 29–31]. In recent years, advances in sequencing and bioinformatics methods have increased the pace of new virus discovery, as one can recover and analyze viral sequences from bulk metagenomes that generated without viral particle enrichment [32]. The untargeted metagenomes may contain sequences of free viruses, integrated proviruses and viruses undergoing the lytic cycle [33], and can reflect the composition of viromes to a large extent. During our study on the microbial communities in the deep-sea hydrothermal vents from Carlsberg Ridge (CR) of Northwestern Indian Ocean, we have also revealed a large number of sequences that were annotated as virus genes, indicating the unexplored viral diversity in these hydrothermal vent fields. Here, we provide the characterization of the community structure, the virus-host interactions and the potential ecological roles of viruses in the CR hydrothermal vents and seven hydrothermal vent sites across the global oceans. So far, this is the largest survey of viruses from the deep-sea hydrothermal vents. A collection of hydrothermal vent viral genomes was recovered from microbial metagenomic datasets and the unique features of vent viral populations were revealed. The results will expand our understanding of viral diversity and functions in the hydrothermal vent ecosystems.

## Results and Discussion

### Overview of the CR hydrothermal vent metagenomes

To explore the community composition and metabolic functions of microorganisms in the CR hydrothermal vents, metagenomes were sequenced from five plume and background seawater samples collected at “Wocan” and “Tianxiu” hydrothermal fields (Table 1). BLAST analysis indicated that about 73% of the predicted genes showed sequence similarity (e-value < 10^−5^) to the NCBI non-redundant (NR) protein database. Of these, most genes had top BLAST hits to bacteria, accounting for 63% and 81% of the mapped metagenomic reads in seawater and plume samples, respectively (Fig. 1). At the phylum level (class level for Proteobacteria), Gammaproteobacteria (on average 22% of the mapped reads), Campylobacteraeota (previously known as Epsilonproteobacteria, 7%), Bacteroidetes (7%), Alphaproteobacteria (7%) and Deltaproteobacteria (5%) were predominant in the hydrothermal environments. A significant difference between seawater and plume samples is the relative abundance of Campylobacteraeota (0.3% versus 13%), a group of chemolithotrophic primary producers who mainly use sulfur compounds and hydrogen as electron donors [34]. The dominant archaeal lineage is Thaumarchaeota, representing 24% and 10% of the seawater and plume metagenomes, respectively. This phylum is particularly abundant in the deep, dark ocean, and has a significant role in biogeochemical cycling of nitrogen and carbon [35, 36]. Sequences derived from eukaryotes account for 5% and 4% of the total mapped reads within seawater and plume datasets, respectively, and fungi of the phylum Ascomycota turn out to be the most abundant eukaryotic group (Fig. 1). Reads representing viral genes also constitute a nonnegligible part of the hydrothermal vent metagenomes (∼2% of the mapped reads in both seawater and plume metagenomes), while genes remaining unclassified account for 1% and 4% of the mapped reads in seawater and plume metagenomes, respectively.

**Figure 1.**
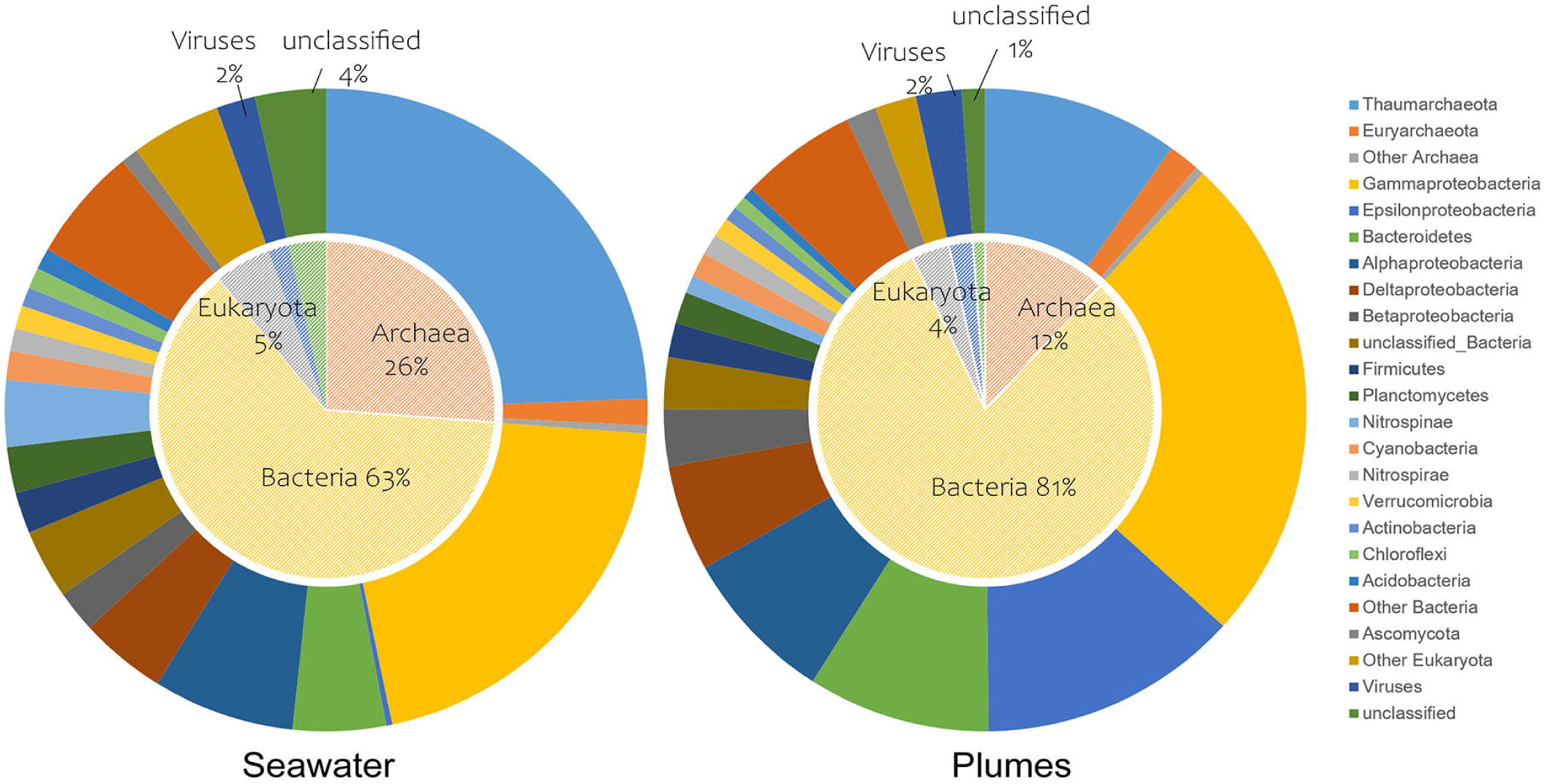
Geographic distribution of metagenome sampling sites involved in this study. The site name, sample type, and sampling depth range in meters below sea level (m) are shown.

**Table 1.**
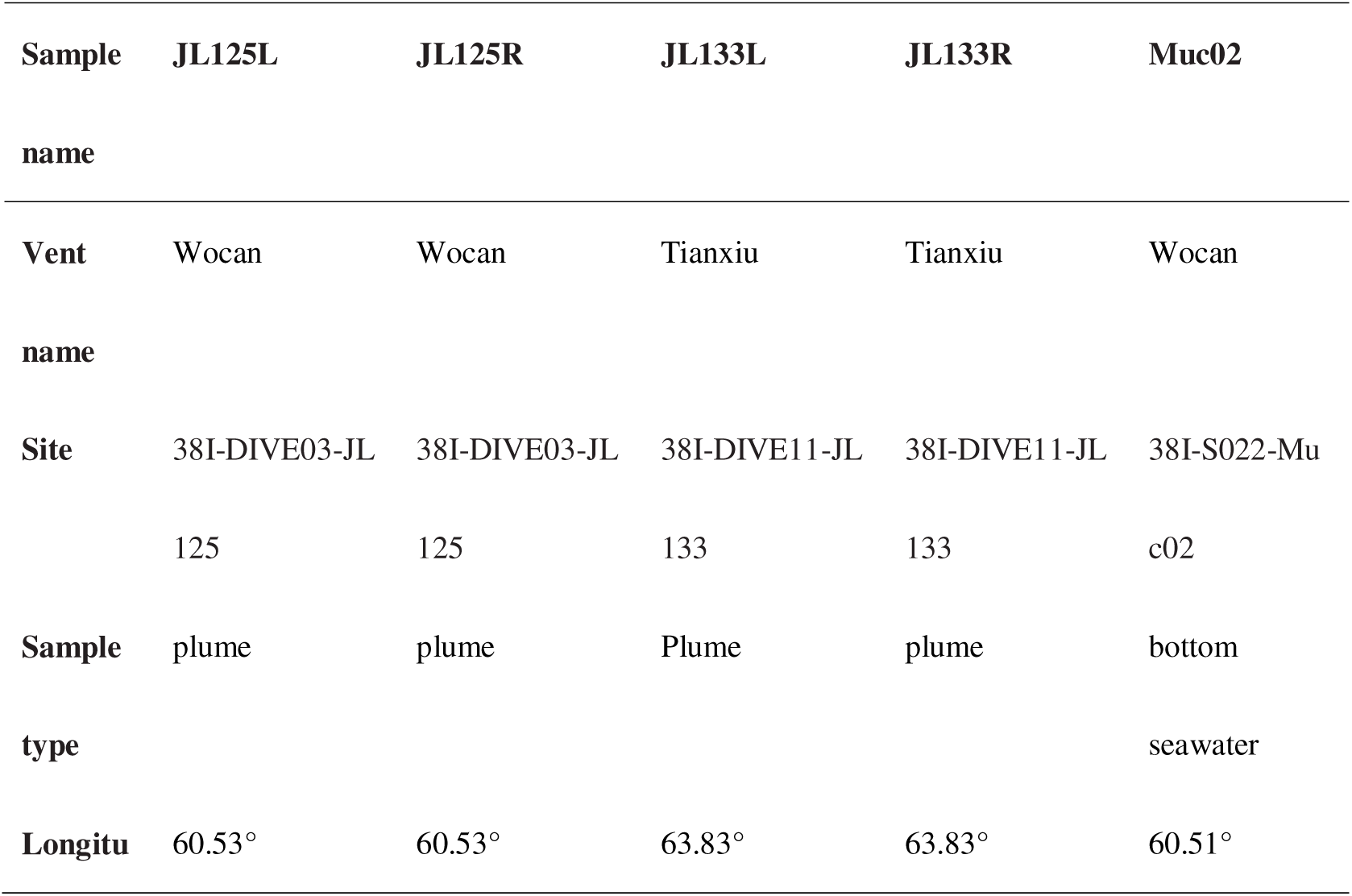
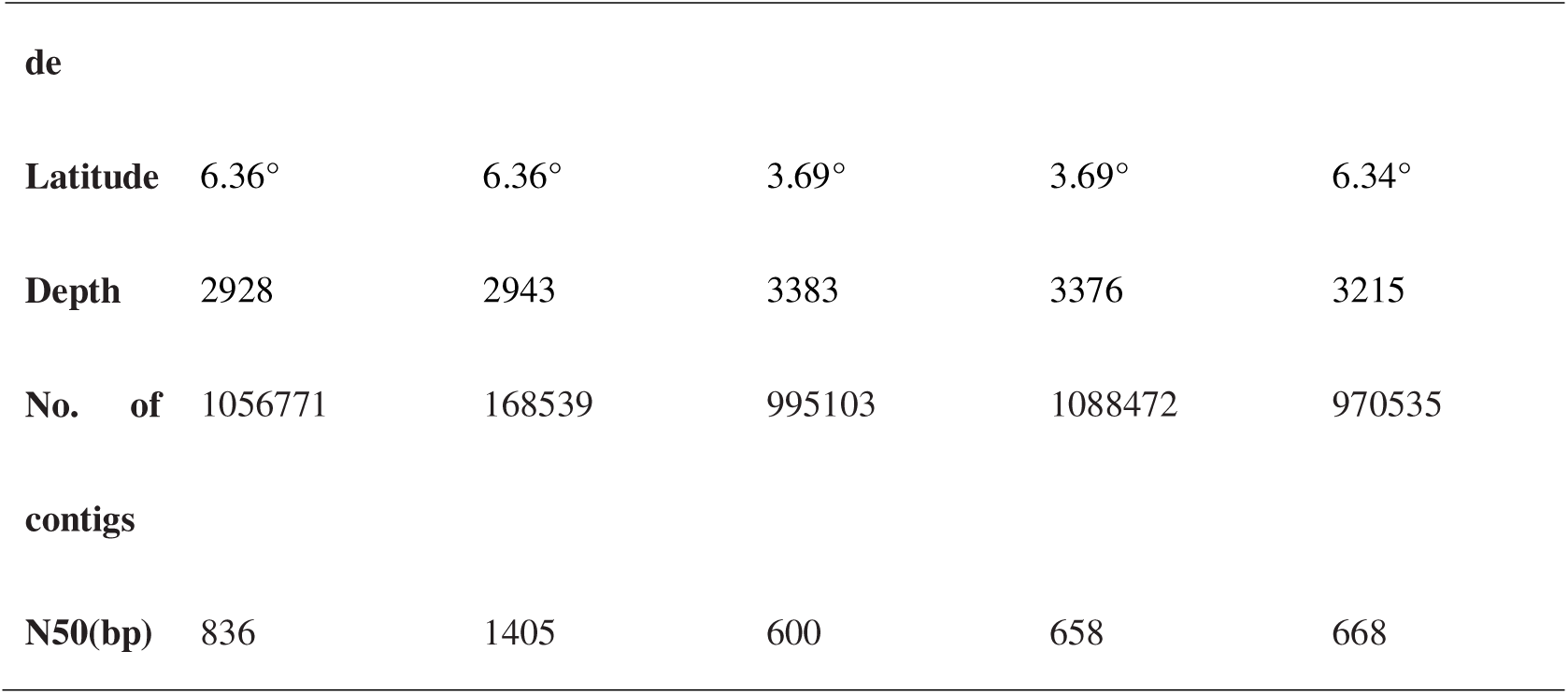

| Sample | JL125L | JL125R | JL133L | JL133R | Muc02 |
| --- | --- | --- | --- | --- | --- |
| name |  |  |  |  |  |
| Vent | Wocan | Wocan | Tianxiu | Tianxiu | Wocan |
| name |  |  |  |  |  |
| Site | 38I-DIVE03-JL | 38I-DIVE03-JL | 38I-DIVE11-JL | 38I-DIVE11-JL | 38I-S022-Mu |
|  | 125 | 125 | 133 | 133 | c02 |
| Sample | plume | plume | Plume | plume | bottom |
| type |  |  |  |  | seawater |
| Longitu | 60.53° | 60.53° | 63.83° | 63.83° | 60.51° |
| **de** |  |  |  |  |  |
| **Latitude** | 6.36° | 6.36° | 3.69° | 3.69° | 6.34° |
| **Depth** | 2928 | 2943 | 3383 | 3376 | 3215 |
| **No. of** | 1056771 | 168539 | 995103 | 1088472 | 970535 |
| **contigs** |  |  |  |  |  |
| **N50(bp)** | 836 | 1405 | 600 | 658 | 668 |

*De novo* assembly and binning of these metagenome sequencing data resulted in the reconstruction of 77 prokaryotic MAGs with ≥70% completeness and ≤10% contamination. These MAGs represent species-level groups that spanning 17 phyla, including 66 bacterial and 11 archaeal MAGs (Supplementary Fig. 1). Most of these MAGs belong to the dominant lineages such as Gammaproteobacteria, Campylobacteraeota, Bacteroidetes and Thaumarchaeota. In addition, 197 high-quality MAGs were recovered from another 29 samples collected at seven different hydrothermal vent sites of Pacific Ocean, Atlantic Ocean and Southwest Indian Ocean (Fig. 2, Supplementary Table 1). These high-quality MAGs and 307 medium-quality MAGs (≥50% completeness and ≤10% contamination) recovered from all vent metagenomes, the publicly available (and some unpublished) genomes of hydrothermal vent microbial isolates, combined with 440 high-quality single-cell amplified genomes (SAGs) retrieved from the CR hydrothermal vents, provide a good basis for investigating the connections between viruses and prokaryotes.

**Figure 2.**
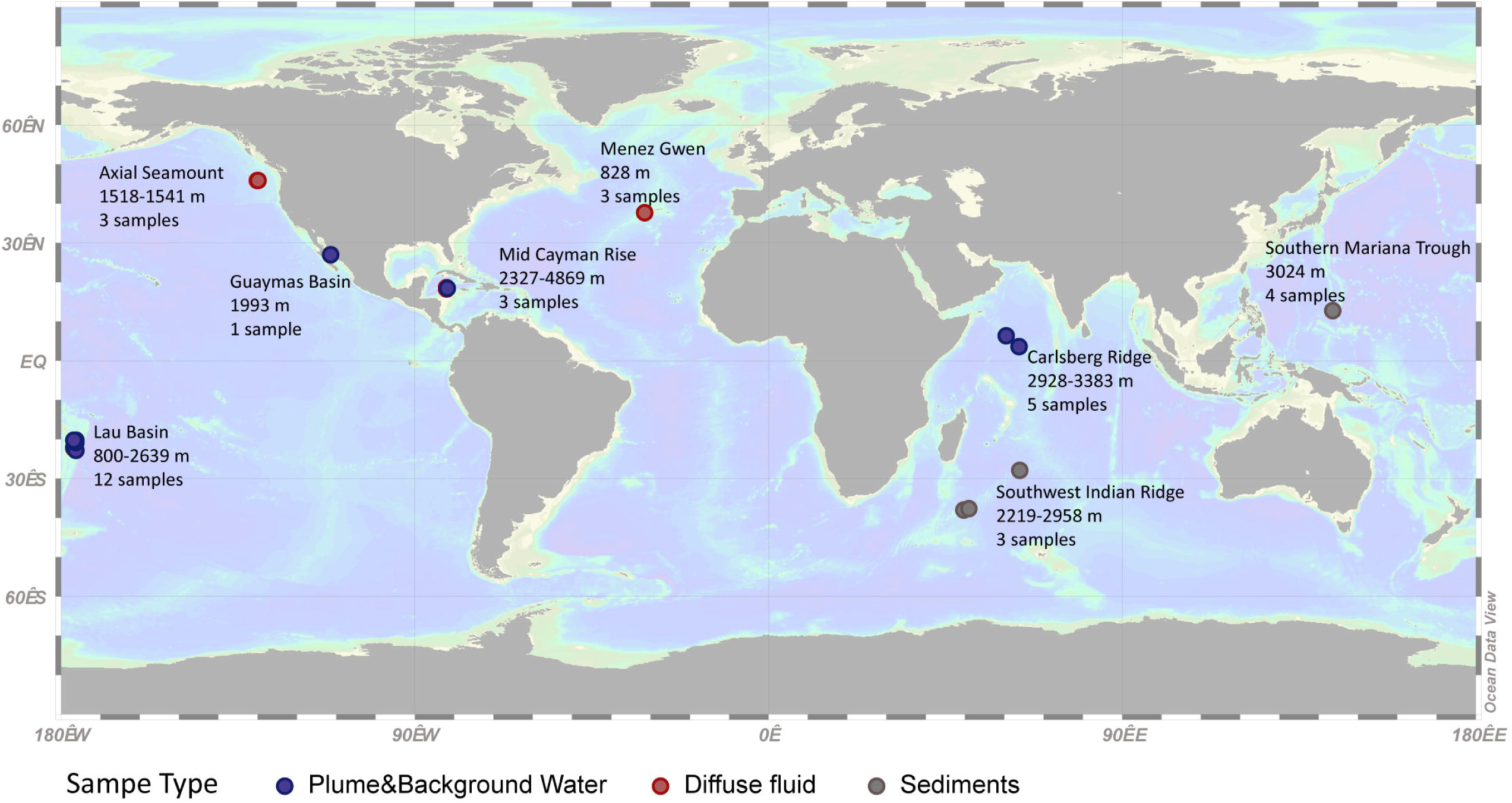
Taxonomic composition of hydrothermal vent plume and background water in the Carlsberg Ridge of Northwestern Indian Ocean. The relative abundance of taxons are calculated at the domain and phylum level (class level for Proteobacteria).

### Diversity and phylogeny of hydrothermal vent viruses

The CR hydrothermal vent metagenomes and additional 29 publicly available metagenomic data sets from seven globally distributed hydrothermal vents were used for virus recovery (Fig. 2). VirSorter v1.0.6 [37] and VIBRANT v1.2.0 [38] were used to identify viral contigs in the hydrothermal vent metagenomes, followed by manual curation. Contigs ≥ 5 kb or ≥ 2kb and circular were pooled together, resulting in 8,847 putative viral sequences. We have retained the small circular contigs because these may represent small single-stranded DNA (ssDNA) virus genomes (such as phages of the family *Microviridae*, and eukaryotic viruses of the family *Circoviridae, Geminiviridae* and *Smacoviridae*, etc.). The 8,847 candidate viral contigs were then clustered at 95% ANI over 80% of the sequence length, producing 4,662 non-redundant vOTUs (Supplementary Table 2). These vOTUs ranged from 2,000 bp to 226,341 bp in size (contig total length= 37,751,900 bp, N50=10,916 bp), including a vOTU larger than 200 kb, which could be classified as “huge phage” [39]. As determined by CheckV [40], almost half of the vOTUs (47%) represented viral genomes of medium quality and above (Supplementary Fig. 2A). Based on the presence of terminal repeats and provirus integration sites, 1,731 vOTUs were identified as complete viral genomes (37%). Of these, 1,727 vOTUs were probably circular, free viruses and four were predicted to be intact proviruses.

A large proportion of the hydrothermal vent vOTUs were classified as double stranded DNA (dsDNA) viruses of the order *Caudovirales* (45%), of which the families *Myoviridae* and *Siphoviridae* were predominant (Supplementary Fig. 2B). As the terminase large subunit (TerL) gene is conserved in all head-tail phages [41], phylogenetic analysis of TerL gene was performed to assess the diversity and genetic distance of *Caudovirales* in hydrothermal vents. Within the viral contigs, a total of 638 complete ORFs encoding the TerL genes were identified and used to construct the phylogenetic tree (Fig. 3A). Most of the sequences from hydrothermal vent metagenomes fell into 14 known lineages which were defined by different DNA packaging strategies, while the rest of them formed several novel branches, indicating the remarkable diversity of head-tail phages in our datasets.

**Figure 3.**
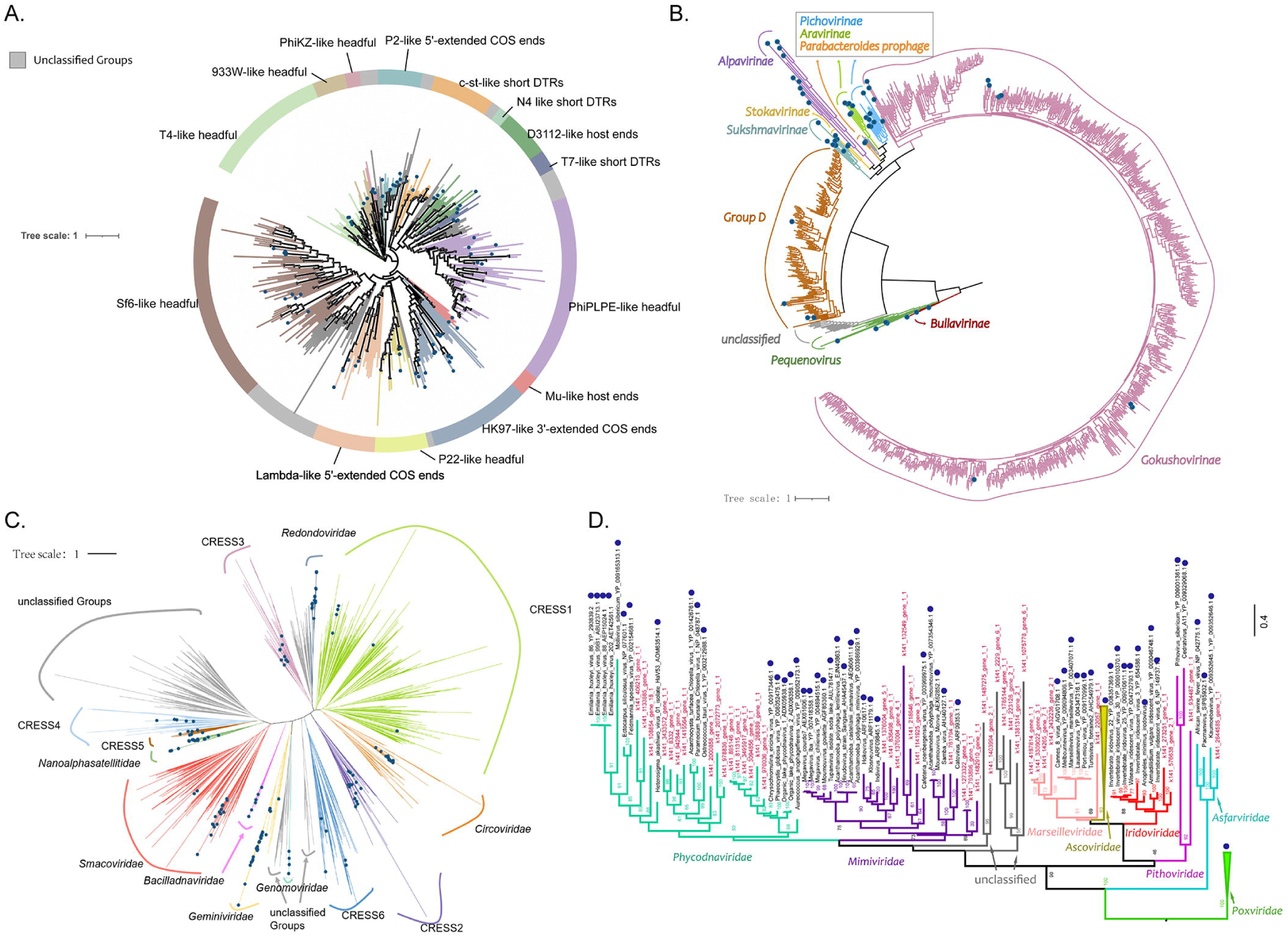
Phylogeny of the major viral groups in hydrothermal vents. Representative members of previously reported virus groups are indicated as blue dots, and potentially novel clades are colored in grey. (A) Maximum likelihood tree of the terminase large subunit protein (terL) of *Caudovirales*. The tree was bootstrapped with 1000 sub-replicates, and bootstrap scores >70% are flagged with dots. (B) Maximum likelihood tree based on VP1 amino acid sequence of Microviridae. (C) Maximum likelihood tree based on Rep proteins of CRESS-DNA viruses. (D) Maximum likelihood tree based on MCP sequences of NCLDVs.

ssDNA viruses accounted for about 36% of the total set of vOTUs, with primary assignment to the family *Microviridae*. Phages of family *Microviridae* are among the smallest DNA viruses [42], with circular genomes ranging from 4.2 to 6.5 kb [43, 44]. A total of 872 complete or near-complete genomes of *Microviridae* were recovered in hydrothermal vent metagenomes. The amino acid sequences of the well-conserved major capsid protein VP1 were used as a phylogenetic marker for classification of these viruses. Majority of the VP1 sequences showed <70% identities to their best matches in NCBI’s NR database and showed <70% identities to each other, reflecting high levels of divergence. Phylogenetic analysis showed that most (694 genomes) of the hydrothermal vent-derived *Microviridae* belonged to the subfamily *Gokushovirinae*, followed by the Group D (143 genomes). A few sequences were clustered within the clades of *Pequenovirus* (8 genomes)*, Pichovirinae* (4 genomes), *Bullavirinae* (1 genome) and *Alpavirinae* (1 genome), and the other 22 genomes were not clustered with any known subfamilies (Fig. 3B). Interestingly, the new clade seemed to consist of viruses with small genomes (around or less than 4 kb). One of these has a genome size of 3,559 bp and encode only three putative ORFs, including a capsid protein, a replication initiator and a protein of unknown function. It represents the smallest microvirus recovered in our datasets, and also the smallest ssDNA phage with the least number of ORFs reported to date.

CRESS-DNA (Circular Rep Encoding Single-Stranded DNA) viruses including *Circoviridae* and its related families were also highly represented (15% of the vOTUs, Supplementary Figure 2B, Supplementary Table 2). This group of eukaryotic viruses have small circular genomes and commonly encode only 2 proteins, of which the replication initiation protein (Rep) is the only universally conserved gene [45]. Based on the sequences of Rep proteins, 694 CRESS-DNA viruses identified from hydrothermal vent metagenomes were clustered within 13 known families, while the other 145 sequences defined several potentially new clades (Fig. 3C).

The nucleocytoplasmic large DNA virus (NCLDVs) is another group of eukaryotic viruses [46], including the families *Poxviridae*, *Iridoviridae*, *Ascoviridae*, *Asfarviridae*, *Marseilleviridae*, *Mimiviridae* and *Phycodnaviridae*, as well as several lineages of unclassified viruses. Viruses of this group were also present in our hydrothermal vent metagenomics datasets (Supplementary Figure 2B, Supplementary Table 2). As shown in Supplementary Table 2, 27 vOTUs were classified as viruses from the *Mimiviridae, Phycodnaviridae* or other NCLDV families. However, the sizes of these vOTUs ranged from 6,112 to 24,611 bp, and only represented small genome fragments of the viruses. Besides, about 19% of the total vOTUs did not show any significant sequence similarity to any known viral families and could not be taxonomically classified for the time (Fig. 3D).

It should be noted that all the hydrothermal plume, diffuse fluid and background seawater samples used in this study were passed through the 0.22 μm filters. Giant viruses, integrated proviruses and actively infecting viruses within the cells would be retained on the membrane, while most free virus particles of small size would be lost in this step, such as the non-tailed ssDNA viruses. Thus, we analyzed the metagenomes of the cellular fraction and the virus-like particle (VLP) fraction of the sediment samples from Southwest India Ridge [19] to evaluate the extent to which they reflect the viral diversity. The results showed that the cellular fraction was comparable to the VLP fraction in aspect of viral recovery (data not shown). The number of recovered CRESS-DNA viruses in VLP fractions doubled those recovered in cellular fractions, but the number of microviruses recovered in cellular fractions was more than those recovered in VLP fractions, despite their small size. One explanation for the differences is that the intracellular microviruses were captured on filters while most of the CRESS-DNA viruses replicating in eukaryotic hosts were excluded from sampling. No significant difference was observed for other viral groups. Therefore, we supposed that the viral sequences identified in the cellular metagenomes included both lytic and lysogenic viruses and could represent the viral diversity to a large extent, at least for the prokaryotic viruses. In this study, we focused on the cellular fractions to compare the viral diversity and distribution patterns across different vent sites.

### Viral communities across different zones of hydrothermal vents

To investigate the viral community structure in hydrothermal ecosystems, the relative abundances of vOTUs in each metagenome were calculated and normalized (Fig. 4, Supplementary Fig. 3). The 34 samples used in this study were collected from 8 different hydrothermal vents across various geographical zones, including hydrothermal plume, background water, diffuse fluid and sediment samples. As a result, the vOTU abundance patterns were primarily clustered by sample types and secondarily by hydrothermal vent sites (Supplementary Fig. 3). The viral community composition of hydrothermal vent sediments were significantly different from those of other habitats. Hydrothermal plumes and the surrounding deep-sea water samples showed similar vOTU abundance patterns, as plume and water samples from the same hydrothermal vent field were always clustered together. This is in agreement with previous studies that suggested the plume microbial communities resemble those from background seawater samples [5], indicating that hydrothermal plumes are strongly influenced by the ambient seawater. Similar to other studies of marine viromes, the majority of the hydrothermal vent viral communities are composed of unclassified viruses (up to 69% of the total viral reads, Fig. 4). At the family level, the tailed phage *Myoviridae* (on average 9.2%, 7.8% and 1.5% in plume, fluid and sediment samples, respectively) was the most dominant group of dsDNA viruses in most samples, followed by *Siphoviridae* (9.1%, 4.6% and 2.4%).

**Figure 4.**
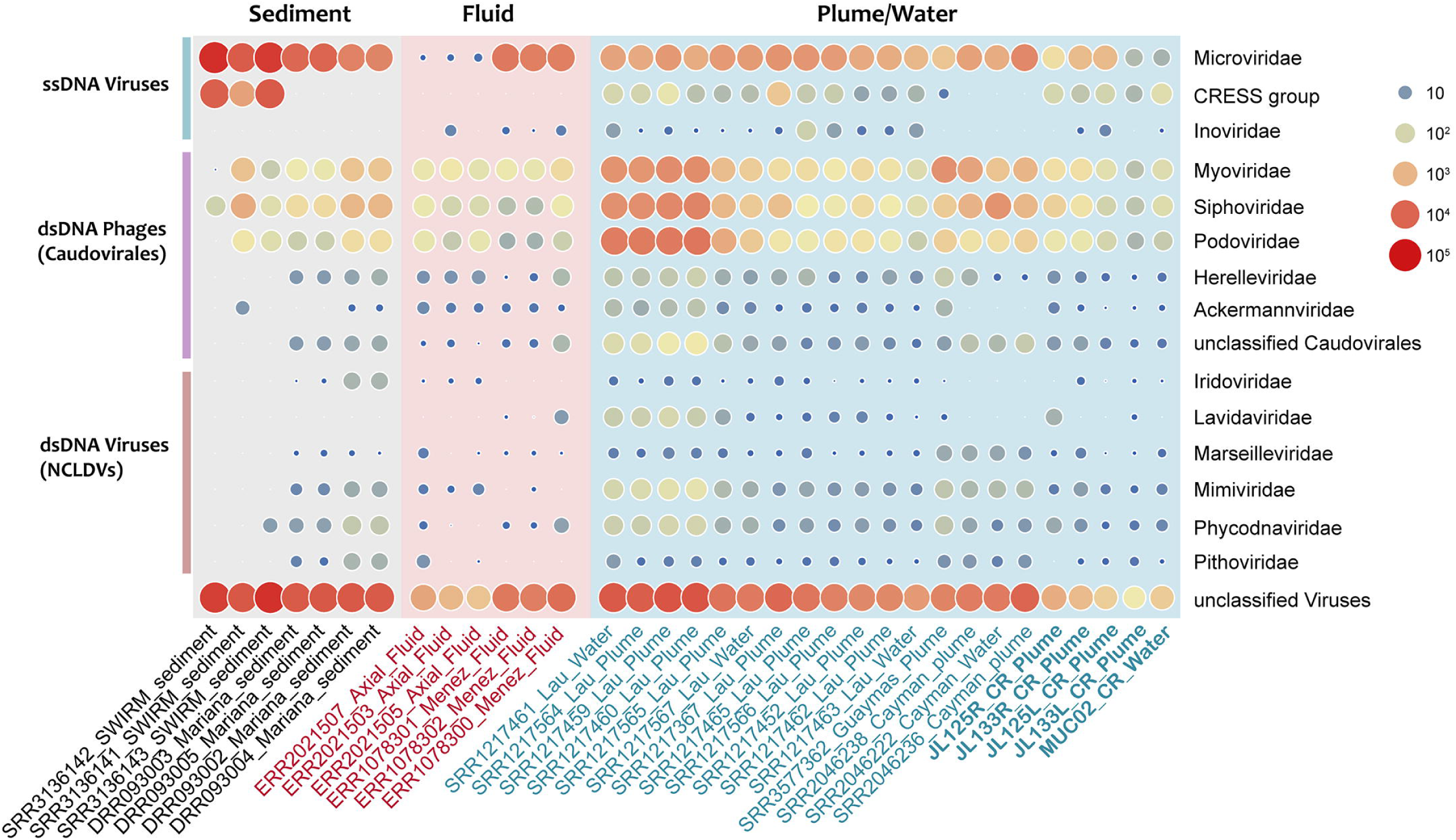
Bubble plot of the relative abundance of viral groups in different hydrothermal samples. The metagenomic samples from this study are indicated in bold.

The ssDNA viruses also accounted for a large fraction of viral communities. In all of the sediment metagenomes, the relative abundance of *Microviridae* were higher than any other dsDNA virus families. The CRESS-DNA virus group were present in most plume samples and three of the sediment samples, accounting for about 2.2% of the total viral reads (Fig. 4). It is possible that the enrichment of ssDNA viruses in these datasets were caused by the multiple displacement amplification (MDA) process, which used the phi29 DNA polymerase and preferentially amplified small circular ssDNA molecules [47, 48]. However, quantification of viral DNA without amplification also revealed the dominance of ssDNA viruses in the total DNA viral assemblages of deep-sea sediments [49]. Thus, we believe that ssDNA viruses are abundant and playing an important role in the hydrothermal vent environments.

The eukaryotic NCLDVs accounted for 0.4% of the total viral reads in hydrothermal plume samples, on average. Of these families, the *Mimiviridae* and *Phycodnaviridae* are the most abundant. In contrast, samples from diffuse fluid and sediment contained less NCLDVs. The large DNA viruses of this group have been frequently detected in marine metagenomes, including those derived from hydrothermal vents [29, 50, 51]. NCLDV sequences in metagenomics datasets may come from marine unicellular eukaryotes or free giant viruses, but their roles in these ecosystems are largely unknown.

Overall, these results showed that the virome structures varied across different hydrothermal vent habitats and different hydrothermal vent fields. To gain a further insight into the viral diversity and distribution, we then used an extensively validated, network-based method [52] to investigate the relationship among hydrothermal vent vOTUs and viral sequences identified from other marine ecosystems.

### Hydrothermal vent viruses are novel and endemic

The 4,662 vOTUs recovered from hydrothermal vents were compared to NCBI Viral RefSeq v97 and viral contigs in other marine metagenomic datasets: GOV 2.0 seawater [53] and cold seeps [54]. A gene-sharing network strategy was used to *de novo* predict genus-level groups (viral clusters, VCs) from viral population data using vConTACT2 [52]. Finally, a total of 16,618 clusters were generated, reflecting the huge and unexplored diversity of marine viruses (Fig. 5A).

**Figure 5.**
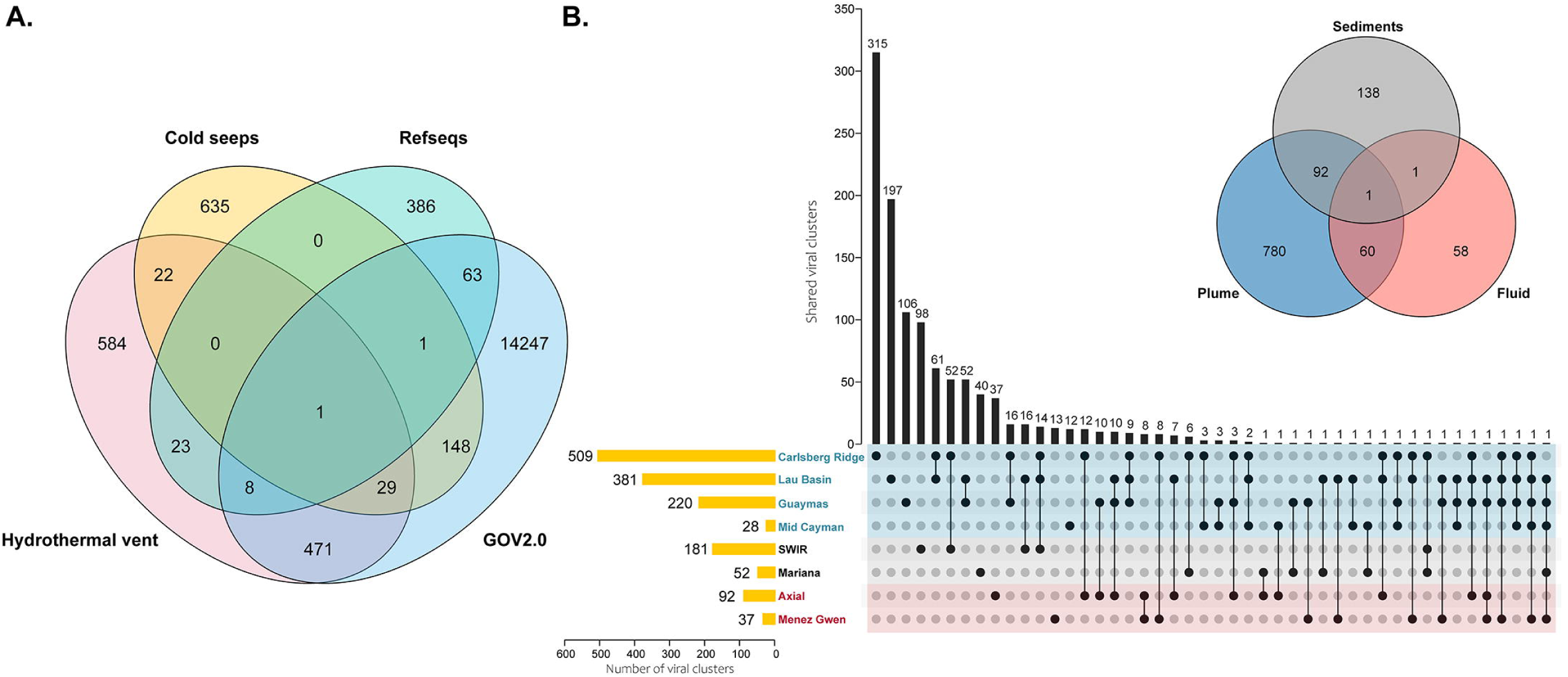
Gene-sharing network analysis of viruses within and beyond hydrothermal vent environments. (A) Venn diagram of shared viral clusters among different environmental virus data sets and RefSeq. (B) Upset plots of shared viral clusters among different hydrothermal vent sites.’

The GOV 2.0 datasets, as the largest marine virus database to date, contributed the largest number of VCs (14,968 clusteers), while taxonomically known viruses from NCBI RefSeq only formed 482 clusters. Consistent with previous studies [54], the viral compositions in different habitats varied considerably, with only 30 clusters shared by all of the three marine metagenomic datasets (Fig. 5A).

The 4,662 hydrothermal vent vOTUs were grouped into 1,138 genus-level clusters, of which only 32 VCs contained genomes of known viruses from NCBI RefSeq. Moreover, **584 VCs (∼51%) were exclusively composed of hydrothermal vent viruses, which may represent completely new candidate genera (Fig. 5A). These VCs contained 1,107 vOTUs, most of which were ssDNA viruses within *Microviridae* (548 vOTUs) and the CRESS-DNA virus families (122).** Around a third of the vOTUs (314) could be classified as tailed phages, probably of novel genera within *Myoviridae* (144), *Podoviridae* (65), *Siphoviridae* (57) and unclassified *Caudovirales* (48). The remaining 123 vOTUs could not be taxonomically classified at the family or even higher levels.

Within hydrothermal vent habitats, a high proportion of the viruses seemed to be endemic, as a majority (818 VCs, 71.9%) of the VCs only occurred in a certain vent site (Fig. 5B). This observation was in congruence with previous work showing that most viruses in hydrothermal vent fluids have limited distributions [31]. Generally, there were more VCs present in hydrothermal plumes than in sediments or diffuse fluids, suggesting a greater viral diversity in hydrothermal plumes. Although the viral communities of different hydrothermal plume samples were somewhat similar at family level, they shared a small fraction of clusters. The number of clusters shared between sediments and fluid were even less, and only one cluster was detected across all sample types (Fig. 5B). The cluster turned out to be viruses of the family *Microviridae*, reinforcing the ubiquity of this group.

### Virus-host connections in hydrothermal vents

The interactions between viruses and their hosts exert a strong influence on the microbial diversity, and are essential for understanding the ecology and functioning of the microbial communities [16]. We sought to link the hydrothermal vent vOTUs to their potential hosts by using a combination of four *in silico* methods [55]. As a result, putative targeted hosts were predicted for a small fraction (494 vOTUs, ∼11%) of the hydrothermal vent vOTUs (Fig. 6, Supplementary Table 3). Specifically, 230 vOTUs were linked to MAGs and SAGs derived from hydrothermal vents. Most connections were predicted by WIsH and CRISPR spacer matching (252 viral-host pairs each), 114 pairs by sequence homology and 119 pairs by tRNA matching. Among them, the linkage of 78 viral-host pairs were supported by two or more prediction strategies. The majority (∼89%) of these vOTUs were predicted to infect a specific host, and only 31 OTUs were linked to hosts from different phyla. This is consistent with the common perceptions and previous findings that most viruses have a narrow host range [37, 54, 55].

**Figure 6.**
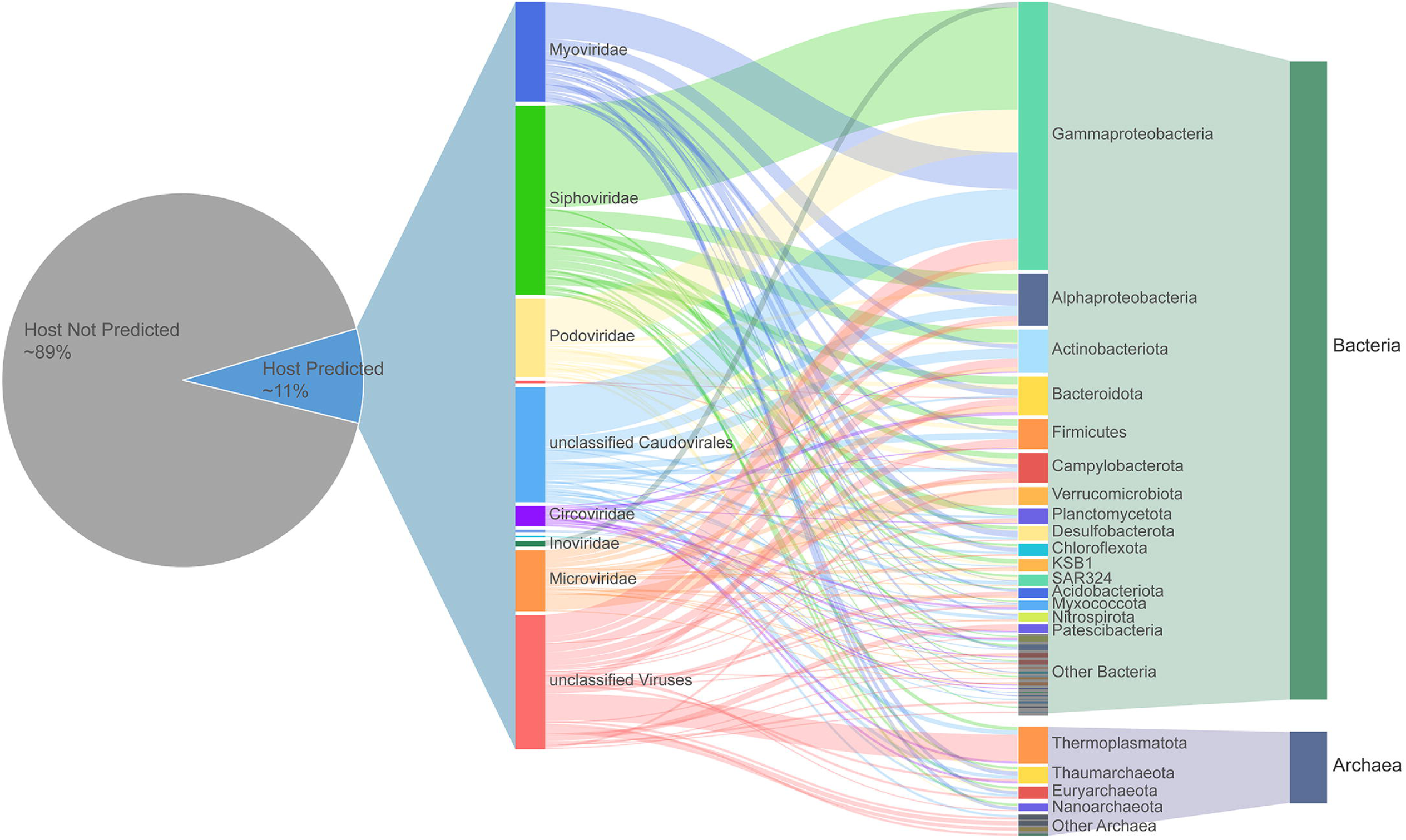
Predicted virus–host linkages in hydrothermal vents.

The predicted hosts of hydrothermal vent phages include bacterial and archaeal species from 39 different phyla (Fig. 6, Supplementary Table 3). Seventy-nine vOTUs were linked to archaea, of which the phylum Thermoplasmatota was the most frequently predicted (33 vOTUs associated).

This is a newly proposed phylum containing the Marine Group II (MGII) and Marine Group III (MGIII) archaea [56]. MGII dominate the ocean surface waters and may play important roles in marine carbon cycle [57], whereas members of MGIII are living in the deep mesopelagic and bathypelagic environments at relatively low abundance [58]. Both these two groups have been found in deep-sea hydrothermal vents and are thought to contribute to organic compounds degradation [59].

Gammaproteobacteria were the most frequently predicted bacterial hosts with 144 vOTUs associated, followed by Actinobacteria (39 vOTUs), Alphaproteobacteria (37 vOTUs), Bacteroidota (34 vOTUs), Firmicutes (27 vOTUs) and Campylobacterota (26 vOTUs). These groups were among the most abundant and active bacterial lineages in the hydrothermal vent ecosystems, as previously reported [7, 10]. For example, Gammaproteobacteria was a large bacterial class with metabolic versatility and was observed in almost all habitats surrounding hydrothermal vents [12]. We found the most frequently predicted hosts within Gammaproteobacteria were the genus *Acinetobacter* (with 15 vOTUs associated), *Alteromonas* (15 vOTUs), *Pseudomonas* (14 vOTUs) and *Alcanivorax* (8 vOTUs), which were dominant in most of the samples involved in this study. According to the well-known Kill-the-Winner hypothesis, abundant microbes are more likely to be infected and lysed by viruses, because high population density will increase host-virus encounter rate [60, 61]. Thus, it is not surprising that plenty of viruses target Gammaproteobacteria in hydrothermal vents. These viruses showed high abundances (Supplementary Fig. 4) and might play important roles in regulating the vent microbial communities.

As another abundant and ubiquitous group that inhabiting the hydrothermal vent environments, Campylobacteraeota were regarded as indicators of hydrothermal activity and as passive tracers of the vent fluid [7, 12, 62]. However, only few potential prophage regions have been reported in the complete genomes of deep-sea Campylobacteraeota [63, 64] and only one of them has been isolated [65] to date. In this study, 26 vOTUs were identified to potentially infect members of the phylum Campylobacteraeota, particularly the genus *Sulfurimonas* (14 vOTUs associated). These included viruses from the family *Myoviridae*, *Podoviridae*, *Siphoviridae, Herelleviridae* and *Microviridae*, while 8 vOTUs remained unclassified at the family level, indicating the unrevealed diversity of viruses infecting hydrothermal vent Campylobacteraeota. Further analysis of these viral genomes will provide new insight into the interactions of this ecologically important group and their phages.

Since the host database we used in this study contains only prokaryotic genomes, it is unlikely to predict the hosts for eukaryotic viruses. However, the results revealed potential connections between some CRESS-DNA viruses and bacterial or archaeal hosts (Fig. 6, Supplementary Table 3). CRESS-DNA viruses from the existing families infect hosts across the eukaryotic domain, including plants, fungi and animals, and the number of CRESS-DNA viruses discovered in metagenomics surveys now far exceeds the number of biologically-characterized viral isolates [45]. These newly identified viruses do not have definitive hosts, but the diversity of the Rep genes implicates that they might have a broader host range than expected [66, 67]. It has been proposed that CRESS-DNA viruses evolved from bacterial rolling circle-replicating plasmids [68, 69]. And recently, a study based on CRISPR analysis suggested that viruses from a CRESS-DNA family infect methanogenic archaea instead of humans [70]. Thus, we could not rule out the possibility that some CRESS-DNA viruses may have hosts beyond eukaryotes.

### The vent viral AMGs involved in various metabolic pathways

Viral infections can affect host metabolism via expression of viral AMGs. To better understand the ecological impact of viruses in the deep-sea hydrothermal ecosystems, we searched AMGs in the hydrothermal vent viral populations and calculated their relative abundances. Based on the comprehensive annotation of viral ORFs, a total of 608 genes were considered to be putative AMGs (Fig. 7, Supplementary Table 4). Sequence homology search against the NCBI’s NR database showed that a large proportion of these AMGs were probably acquired from Proteobacteria, especially the Gamma- and Alphaproteobacteria, while ∼21% of the AMGs were from unclassified source species (Fig. 7B). The origins of AMGs reflected the virus-host connections from the side. According to KEGG annotation, the identified AMGs of hydrothermal vent viruses were involved in a variety of metabolic pathways, including those related with carbohydrate metabolism, amino acid metabolism, and metabolism of cofactors and vitamins (Fig. 7A). This is in consistence with the viral metabolic profiles revealed by analyzing 6 hydrothermal vent metagenomes [38], suggesting that there are some common features in the patterns of metabolic capabilities of hydrothermal vent viromes.

**Figure 7.**
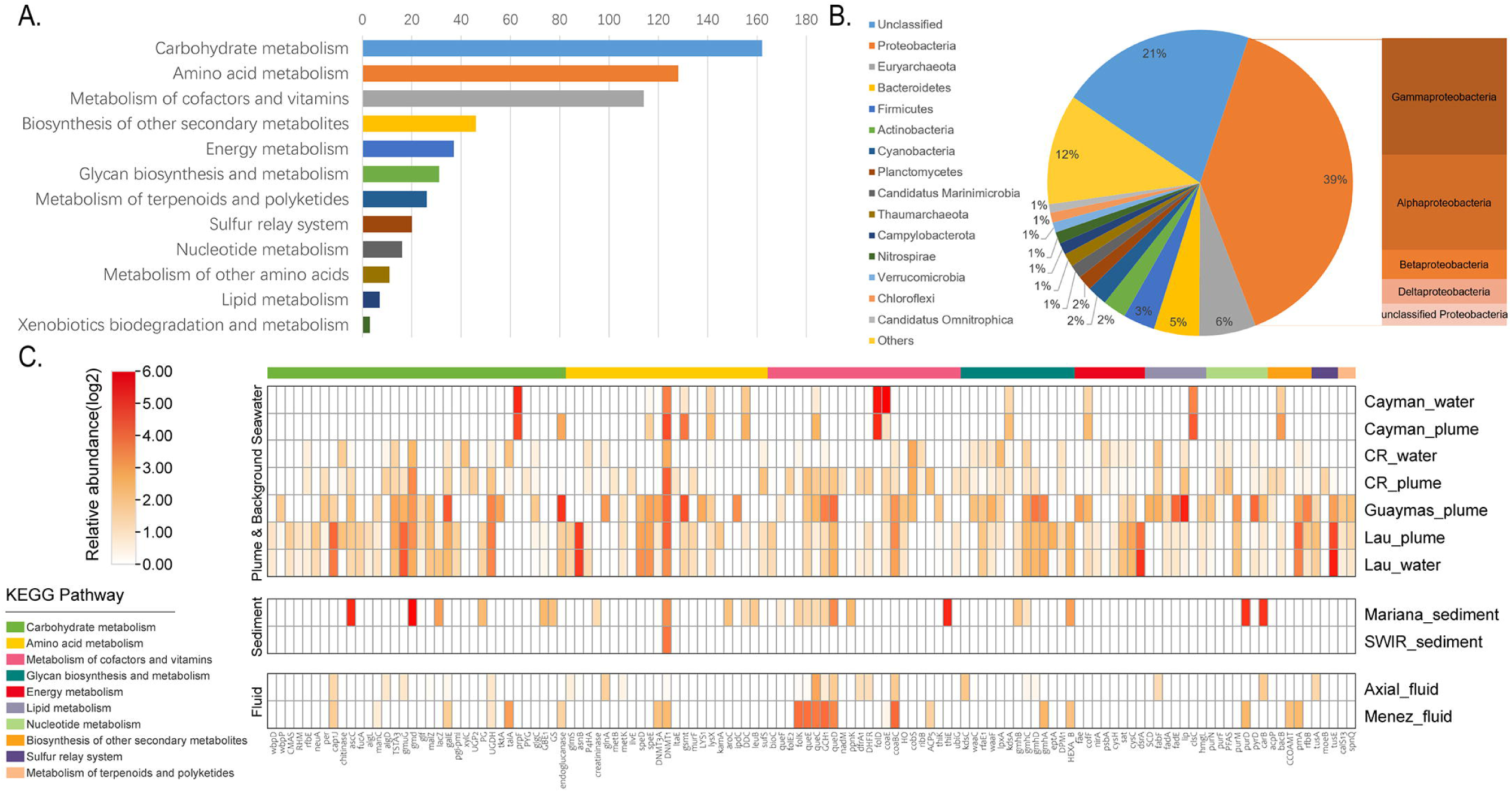
Function and abundance profiles of virus-encoded auxiliary metabolic genes (AMGs). (A) Classification of AMGs into KEGG metabolic categories. (B) Predicted source organisms of viral AMGs. (C) Relative abundance of AMGs in different hydrothermal vent samples.

The AMG composition and abundance profiles across the hydrothermal vent samples indicated that viruses in hydrothermal plumes encoded a larger number of AMGs with diverse functions (Fig. 7C). Besides, most of the AMGs had higher abundances in the plume samples compared with hydrothermal fluid or sediments. This may reflect the role of viruses in facilitating host adaptation to the dynamic nature of hydrothermal plumes, and in congruence with the metabolic versatility of microbes therein [8]. Specifically, AMGs involved in energy metabolism were only found in plume samples, including those related with methane, nitrogen and sulfur metabolisms (Fig. 7C). Several AMGs were associated with the sulfur metabolism pathways, such as genes coding for the phosphoadenosine phosphosulfate reductase (*cysH*), adenylylsulfate kinase (*cysC*), sulfate adenylyltransferase (*sat*) and dissimilatory sulfite reductase subunits A (*dsrA*). The *cysH* and *cysC* genes are involved in assimilatory sulfate reduction, whereas the *sat* and *dsrA* genes are related with dissimilatory sulfur reduction/oxidation [71]. Sulfur oxidation and sulfate reduction are both important parts of the sulfur cycling in hydrothermal vent ecosystems [72], and the presence of these AMGs suggested that phages extensively participate in these pathways.

The most abundant AMG identified in hydrothermal vent metagenomes turned out to be the DNA cytosine methyltransferase (DNMT1, *dcm*). However, viral *dcm* and another 13 AMGs were present in metagenomes derived from several diverse environments and were thus considered to perform central functions in viral life cycle [38]. Except for these globally conserved AMGs, the *tusE* gene related with sulfur relay system was identified in high abundance. This gene encodes a sulfur transfer protein for tRNA thiol modifications, which is required for protein synthesis machinery [73]. Expression of tRNA thiolation genes has been reported to increase the stability of tRNA structure, and was essential for survival at high temperatures in bacteria [74, 75]. In addition, it has been suggested that sulfur relay system was involved in the microbial tolerance against acid stress [76, 77], heavy metal [78] and organic solvents [79]. Therefore, this AMG may benefit the hosts by improving their adaptability to various stress conditions, and thus provide them a growth advantage in the hydrothermal vent environments where temperature and chemistry are fluctuating substantially.

## Conclusions

In this study, we explored the viral community of the CR hydrothermal vents and additional hydrothermal vent sites across a wide geographical area, based on untargeted metagenomic data. The hydrothermal vent viral genome dataset was established for the first time, and the viral distribution, phylogenetic diversity and metabolic potential were characterized as well as their interactions with hosts across different habitats. The viral populations recovered from hydrothermal vents are highly diverse and novel, and most of them tend to be endemic to specific habitats. We also revealed the high abundance and genetic diversity of ssDNA viruses, which are neglected in the previous studies. Many viruses were predicted to infect the ecologically important bacterial and archaeal clades inhabiting the hydrothermal vents, and involved in various host metabolisms including energy conservation and tRNA modification by carrying diverse AMGs, which may benefit their hosts in adverse conditions. Compared to diffuse flow and sediment samples, the viruses in hydrothermal plumes possess a higher diversity and AMG abundance. These results highlight the hidden role of vent viruses and the remarkable impact they exert on vent microbial communities. The virus-host interactions in hydrothermal vents may profoundly influence the biogeochemistry of the deep oceans via vent fluid circulation and plume drift with unique prokaryotic community. With the improvement in deep-sea sampling and culture-dependent and independent technologies, a more comprehensive understanding of virus-host interactions in hydrothermal ecosystems is surely expected.

## Methods

### Sample collection

Plume and background seawater samples were collected from two different deep-sea hydrothermal vents, “Wocan” and “Tianxiu”, at Carlsberg Ridge of Northwest Indian Ocean during the COMRA cruise DY 38 in March 2017 (Table 1). A human-operated vehicle ‘Jiaolong’ was used to collect 1.5-L water samples in individual Niskin bottles. The collected water were filtered through 0.22 μm polycarbonate membranes (diameter 45 mm; Whatman, Clifton, NJ, USA) and were frozen at −80℃ on board for DNA extraction.

### DNA extraction and sequencing

The total DNA was extracted from filtration membranes as described previously [5]. Multiple displacement amplification of genomic DNA was performed using the illustra Ready-To-Go GenomiPhi V3 DNA Amplification Kit (GE Healthcare, Piscataway, NJ, USA). Paired-end library was constructed using NEXTFLEX Rapid DNA-Seq (Bioo Scientific, Austin, TX, USA). Adapters containing the full complement of sequencing primer hybridization sites were ligated to the blunt-end of fragments. Shotgun sequencing was performed on Illumina Hiseq PE150 platform (Illumina Inc., San Diego, CA, USA) at Majorbio Bio-Pharm Technology Co., Ltd. (Shanghai, China) according to the manufacturer’s instructions (www.illumina.com).

### Metagenome assembly and annotation

The raw reads obtained by Illumina pair-end sequencing were trimmed and quality filtered using the fastp software [80]. Clean reads were then assembled using MEGAHIT [81] with default options. Metagene [82] was used to predict protein coding sequences (CDS) from the metagenomic assemblies. Non-redundant genes generated by CD-HIT clustering (with a 95% sequence identity and 90% coverage) were aligned to NCBI NR database using BLASTp (with e-value cutoff of 1e-5) for taxonomic classification. Functional annotations were conducted based on comparisons with KEGG [83], eggNOG v5.0 [84] and the Carbohydrate-Active enZYmes (CAZy) databases [85] using BLASTp [86] program (with e-value cutoff of 1e-5). Additional 29 publically available metagenomic datasets generated from hydrothermal vent samples (Supplementary Table 1) were downloaded from the National Center for Biotechnology Information (NCBI) Sequence Read Archive (SRA) database, and were quality-controlled, assembled and annotated as described above. The map of the sampling locations was created in Ocean Data View v5.5.2 [87].

### Metagenomic binning and metagenome-assembled genome (MAG) classification

Contigs larger than 1500 bp from the final assemblies were included for metagenomic binning using MetaBAT2 v2.12.1 [88] with default parameters. The original bins were then run through the MetaWRAP’s reassemble_bins module [89] to improve their quality, and the completeness and contamination of the resulting bins were evaluated by CheckM [90]. The high-quality bins (completeness ≥ 70% and contamination ≤10%) were then dereplicated at 95% average nucleotide identity (ANI) using dRep v2.3.2 [91], resulting in 77 species-level MAGs. Taxanomic assignment of the MAGs was performed using the GTDB-Tk package v0.3.2 [92], and the phylogenomic relationship of MAGs were inferred using PhyloPhlAn 3.0 based on 400 universal marker genes [93]. Support for nodes in the ML trees was evaluated with 1000 ultrafast bootstrap replicates, and the generated tree was visualized by iTOL v4 [94].

### Identification of viral contigs

Contigs ≥ 2 kb from the metagenome assemblies were used to recover viral sequences. VirSorter analysis [37] was run with the parameter “--db 2 (viromes database)”, and only the highest confidence contig categories 1, 2, 4, and 5 were included in this study, with categories 4 and 5 being manually curated. Contigs containing at least one of the viral hallmark genes (such as ‘virion structure’, ‘capsid’, ‘portal’, ‘head’, ‘tail’, ‘baseplate’ or ‘terminase’) were retained. VIBRANT v1.2.1 [38] was also used to identify viral contigs using the default parameters, and only the complete circular, high- and medium- quality drafts were kept for further analysis. The contigs identified by Virsorter and VIBRANT were then compiled and clustered at 95% nucleotide identity and 80% coverage [95], producing 4,662 viral populations, or viral operational taxonomic unit (vOTUs). Finally, the CheckV pipeline was used to estimate the completeness of the viral genomes and to predict viral lifestyles [40].

### Abundance Profiling in Metagenomics Data

To calculate the relative abundances of viral populations and host microorganisms in each sample, clean reads from metagenomes were mapped to the viral contigs or microbial genomes using the CoverM package (https://github.com/wwood/CoverM) with contig mode and genome mode, respectively. RPKM (reads per kilobase per million mapped reads) values was selected to represent the relative abundances of viral and host populations. Pearson correlation was used to calculate the distances between samples for hierarchical clustering. Heatmaps were generated using the pheatmap R package and the TBtools [96].

### Host prediction

Four computational host prediction strategies were used to identify potential virus-host interactions [55]. (i) CRISPR spacers match. A Clustered Regularly Interspaced Short Palindromic Repeats (CRISPRs) spacer database was created for a set of microbial genomes using the MinCED tool [97]. For metagenomics reads, crass v1.0.1 [98] was used with default parameters to recover CRISPR spacers and repeat elements. The identified spacers were queried for exact sequence matches against all viral contigs using the BLASTn-short mode in the BLAST+ package [99]. Match requirements were at least 95% identity over 95% spacer length, and only ≤1 mismatch was allowed. The corresponding CRISPR direct repeat types were connected to microbial genomes via BLASTn (with e-value cutoff of 1e-10, 100% nucleotide identity) [33]. (ii) Transfer RNA (tRNA) match. tRNAs were recovered from microbial genomes and viral contigs by ARAGORN with “-t” option [100]. The identified tRNA sequences were compared using BLASTn and only a perfect match (100% coverage and 100% identity) was considered indicative of putative host-virus pairs. (iii) Nucleotide sequence homology search [101]. To link prophages with hosts, viral contigs were searched against microbial genomes using BLASTn with the following thresholds: 75% minimum coverage of viral contig length, 70% minimum nucleotide identity, 50 minimum bit score, and 0.001 maximum e-value. (iv) k-mer frequencies. WIsH v1.0 [102] was run with default parameters against the host database. Connections were inferred when pC<C0.001. If multiple hosts were predicted for a vOTU, the one that supported by different approaches was chosen as the most confident. The host database employed for these prediction methods was composed of (i) all reference genomes from the Genome Taxonomy Database (GTDB), (ii) all MAGs (≥50% completeness and ≤10% contamination) recovered from hydrothermal vent metagenomes (n=581), (iii) all SAGs obtained from the CR hydrothermal vent (n=440), and (iv) a custom collections of marine microbial genomes.

### Viral taxonomic assignment and network analysis

The predicted open reading frames (ORFs) of the viral contigs were mapped against the NR protein database using DIAMOND v0.9.21 [103], and their taxonomic affiliations were determined using the CAT v5.0.3 package [104] based on the Last Common Ancestor (LCA) algorithm. Contig classification is based on a voting approach of all classified ORFs by summing all bit-scores from ORFs supporting a certain classification. Protein-sharing network analysis of the hydrothermal vent vOTUs, the reference phage genomes (from NCBI Viral RefSeq version 97), the Global Oceans Viromes 2 (GOV 2.0) datasets [53], and viral contigs from cold seeps [54] was performed by vConTACT v2.0 [52]. Briefly, Prodigal v2.6.3 [105] was used for ORF prediction from the vOTUs. The predicted protein sequences were then subjected to all-to-all BLASTp using DIAMOND, and the BLAST result file were used as input for vConTACT2. The similarity score between vOTUs was calculated based on the number of shared protein clusters, and related vOTUs with a similarity score of ≥1 were grouped into viral clusters.

### Construction of phylogenetic trees

For phylogenetic trees, the deduced amino acid sequences of selected marker genes were aligned using the MUSCLE program [106], and the multiple alignments were trimmed by TrimAl v1.2 [107]. IQ-TREE2 [108] was used to infer the Maximum likelihood (ML) tree with the best substitution model selected by ModelFinder [109], and support for nodes in the ML trees was evaluated with 1000 ultrafast bootstrap replicates. The resulting trees was visualized and annotated using FigTree v1.4.4 (http://tree.bio.ed.ac.uk/software/figtree/) or the iTOL v4 online tool [94].

### Identification of auxiliary metabolic genes

Functional annotation for the ORFs in the viral contigs were conducted based on comparisons with the eggNOG v5.0 [84] database using eggNOG-mapper v2 [110]. Genes with a KEGG annotation falling under the “metabolic pathways” category or “sulfur relay system” were considered to be putative vAMGs, as previously defined in VIBRANT [38]. Finally, manual curation was performed to remove the metabolic genes involved in common viral functions. To generate the abundance profiles for AMGs, clean reads were mapped to the metagenomes using bowtie2 [111], and RPKM values of each gene was calculated. The sum of RPKM values of genes with the same KO annotations was used to represent the relative abundance of each gene category.

## Declarations

### Ethics approval and consent to participate

Not applicable

### Consent for publication

Not applicable

### Availability of data and material

The metagenomic datasets used for analysis in this study are publicly available in the NCBI SRA repository at https://www.ncbi.nlm.nih.gov/sra, and accession numbers are listed in the Supplementary Table 1. Metagenomic data from CR hydrothermal vent were also deposited in the SRA database (accession number PRJNA685608). The sequences of the vOTUs generated from the current study have been deposited in the National Omics Data Encyclopedia (NODE) database at https://www.biosino.org/, accession number OEP003015.

## Funding

This work was funded by the National Key Research and Development Program of China (No.2018YFC0310705; No.2018YFC0310701); the grant of Laboratory for Marine Biology and Biotechnology, Pilot National Laboratory for Marine Science and Technology (Qingdao) (No. OF2019NO05); Scientific Research Foundation of Third Institute of Oceanography, MNR (No. 2018022); National Natural Science Foundation of China (No. 42006088); Natural Science Foundation of Fujian Province of China (No. 2019J05149); and the China Ocean Mineral Resources R&D Association (COMRA) program (No. DY135-B2-01).

### Authors’ contributions

Conceptualization, RC and ZS; methodology, RC and XL; investigation, LJ; data curation, XL, LJ and LG; writing—original draft preparation, RC; writing—review and editing, CG and ZS; supervision, ZS; funding acquisition, ZS and RC. All authors read and approved the final manuscript.

## Acknowledgements

The authors are grateful to the whole R/V Xiang-yang-hong 9 team of the scientific cruise DY 38I. The authors also thank Prof. Rui Zhang and Min Jin for their helpful suggestions and comments on improving the manuscript.

## Supplementary Files

**Supplementary Table 1**. **Metagenome datasets used for analysis in this study.**

**Supplementary Table 2. Characteristics of vOTUs recovered from hydrothermal vent metagenomes.**

**Supplementary Table 3. Predicted virus-host connections for vOTUs.** The most confident host for a vOTU is highlighted.

**Supplementary Table 4. Annotation of putative auxiliary metabolic genes.**

**Supplementary Figure 1. High-quality MAGs recovered from CR hydrothermal vent metagenome.** Maximum-likelihood phylogenetic tree were inferred from 400 universal marker genes using PhyloPhlAn 3.0. MAGs obtained in this study were colored in red, and MAGs that were linked with viruses were indicated by triangle or multiple triangles if more than one linkages were predicted.

**Supplementary Figure 2. Genome quality and taxonomic composition of hydrothermal vent vOTUs.** (A) Proportion of genome quality categories assessed by CheckV. (B) Taxonomic classification of vOTUs at the family level.’

**Supplementary Figure 3. Distribution patterns of all hydrothermal vent vOTUs.** The relative abundances (RPKM values, y-axis) of vOTUs in each sample (x-axis) were displayed on the log2 scale, and are hierarchically clustered by samples.

**Supplementary Figure 4. Distribution patterns of viruses and their predicted hosts in deep-sea hydrothermal vents.** Relative abundances of vOTUs (top left triangle) and their predicted hosts (bottom right triangle) were grouped by the host taxonomy and were displayed on the log2 scale.

