## Supplementary figures and images for "Viruses Diversity and Interactions with Hosts in Deep-Sea Hydrothermal Vents"

### Supplementary Fig. 1

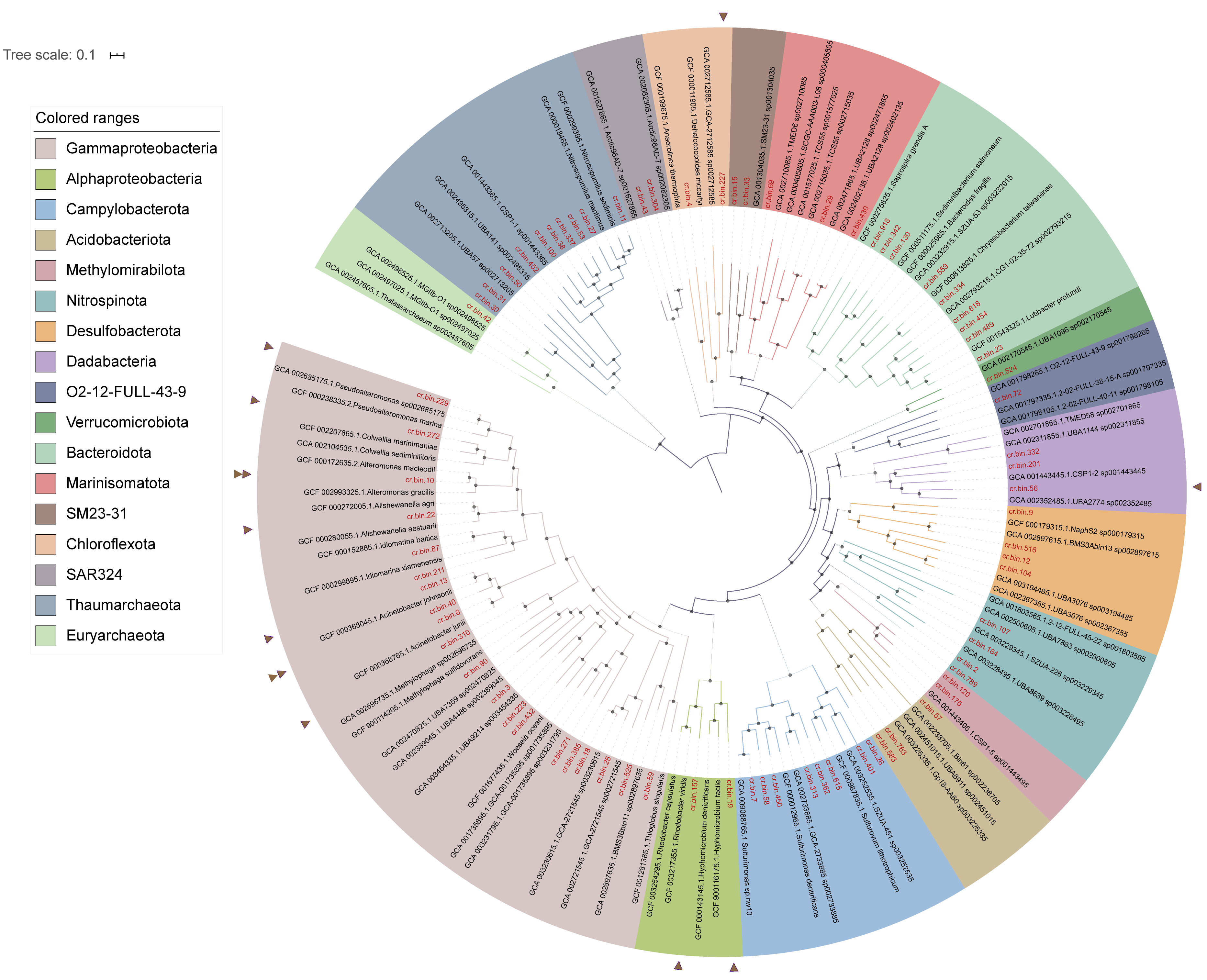

### Supplementary Fig. 2

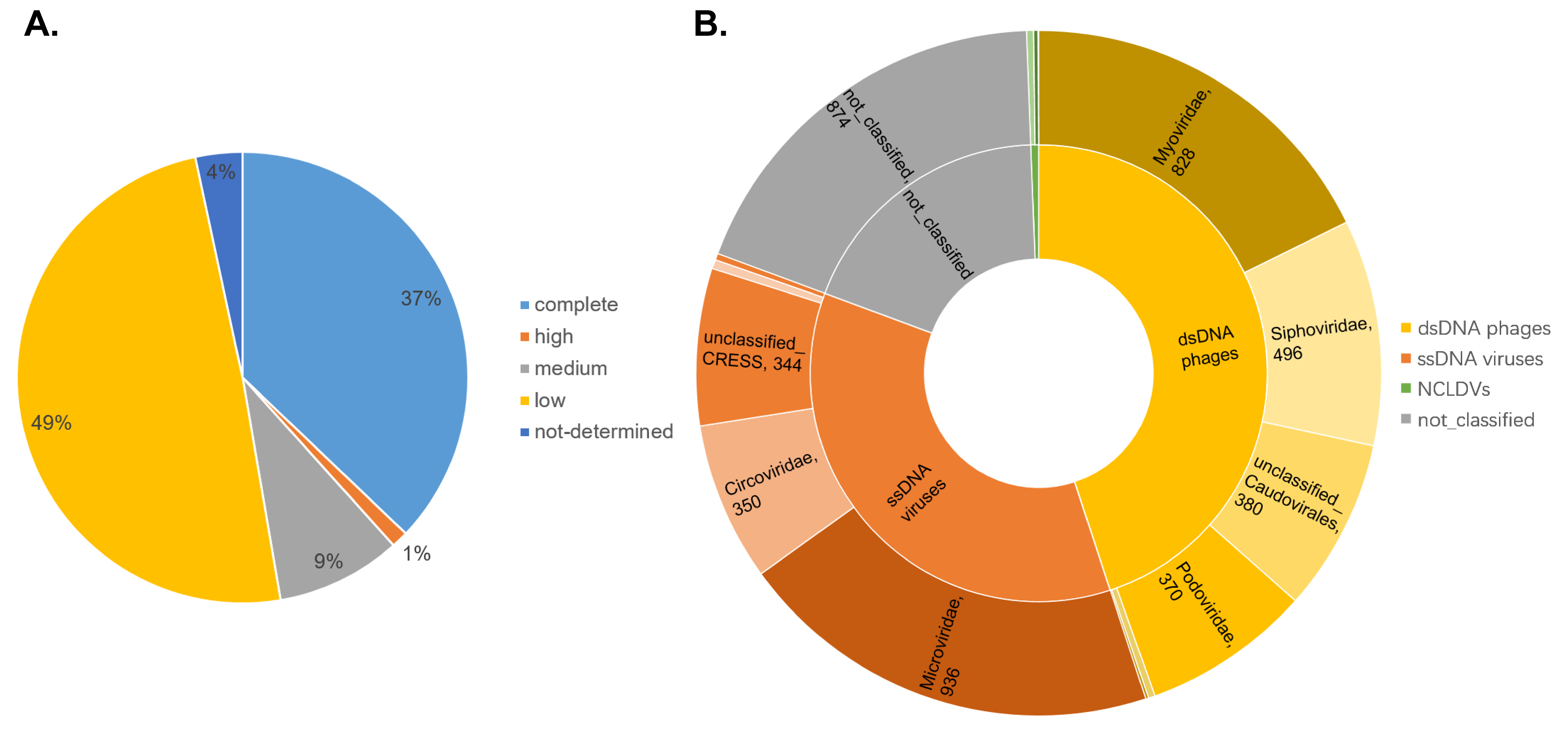

### Supplementary Fig. 3

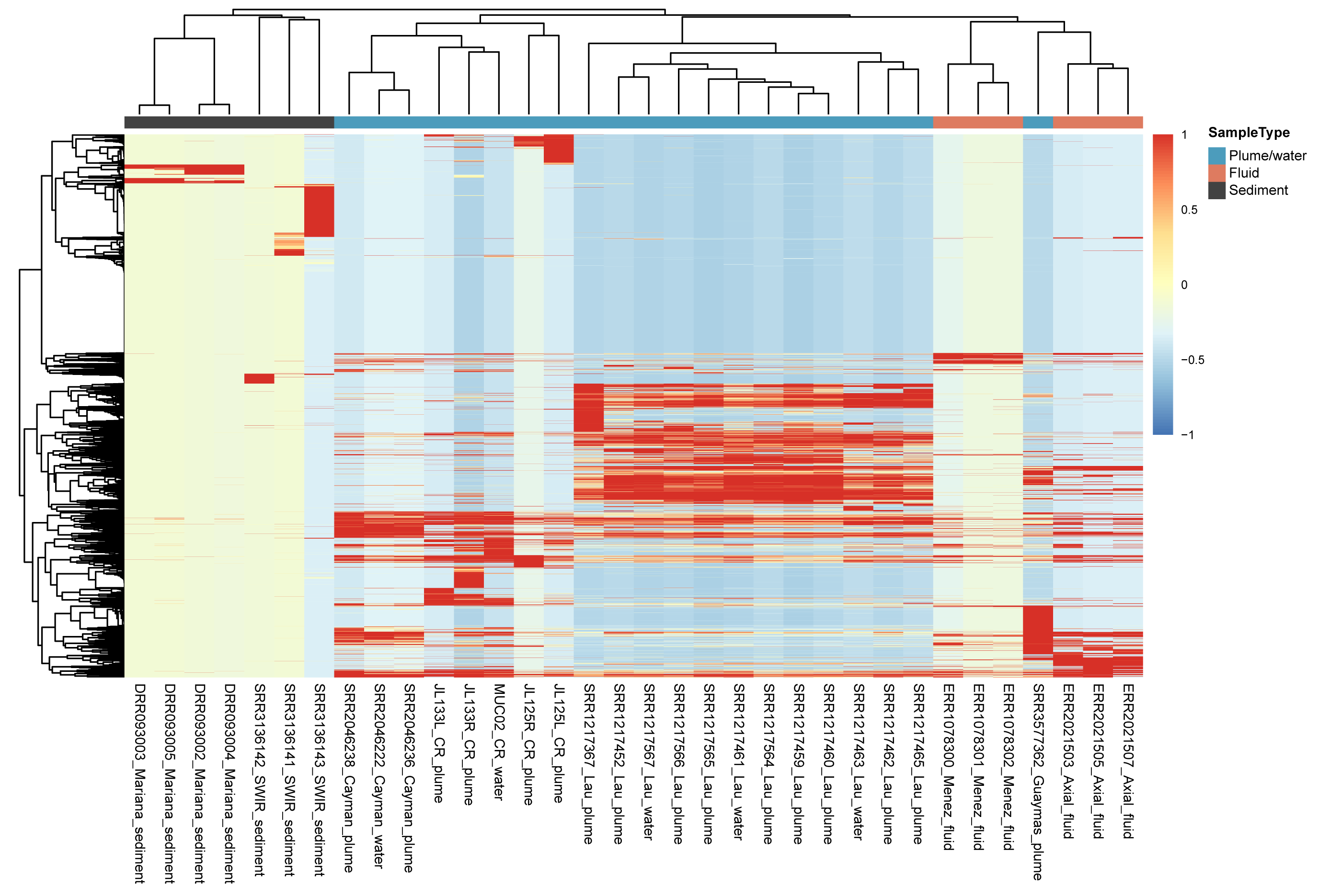

### Supplementary Fig. 4

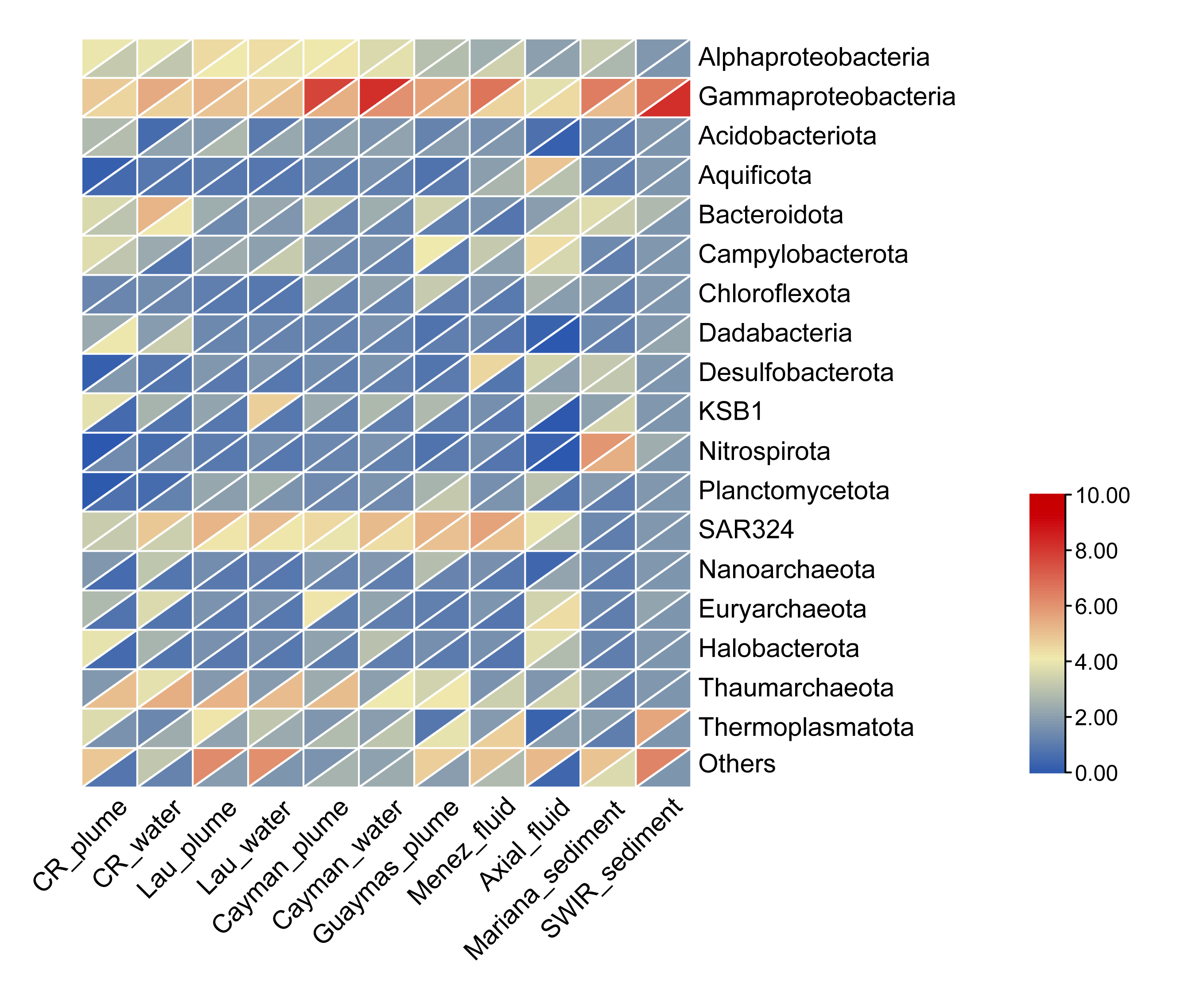
